# The Automated Preprocessing Pipe-Line for the Estimation of Scale-wise Entropy from EEG Data (APPLESEED): Development and validation for use in pediatric populations

**DOI:** 10.1101/2021.07.10.450198

**Authors:** Meghan H. Puglia, Jacqueline S. Slobin, Cabell L. Williams

**Affiliations:** University of Virginia

**Keywords:** multiscale entropy, pediatric EEG, preprocessing pipeline, infant development

## Abstract

It is increasingly understood that moment-to-moment brain signal variability – traditionally modeled out of analyses as mere “noise” – serves a valuable function role and captures properties of brain function related to development, cognitive processing, and psychopathology. Multiscale entropy (MSE) – a measure of signal irregularity across temporal scales – is an increasingly popular analytic technique in human neuroscience. MSE provides insight into the time-structure and (non)linearity of fluctuations in neural activity and network dynamics, capturing the brain’s moment-to-moment complexity as it operates on multiple time scales. MSE is emerging as a powerful predictor of developmental processes and outcomes. However, differences in data preprocessing and MSE computation make it challenging to compare results across studies. Here, we (1) provide an introduction to MSE for developmental researchers, (2) demonstrate the effect of preprocessing procedures on scale-wise entropy estimates, and (3) establish a standardized EEG preprocessing and entropy estimation pipeline that generates scale-wise entropy estimates that are reliable and capable of differentiating developmental stages and cognitive states. This novel pipeline – the Automated Preprocessing Pipe-Line for the Estimation of Scale-wise Entropy from EEG Data (APPLESEED) is fully automated, customizable, and freely available for download from https://github.com/mhpuglia/APPLESEED. The dataset used herein to develop and validate the pipeline is available for download from https://openneuro.org/datasets/ds003710.

## Introduction

Development signifies a time of great complexity and dynamism. Changes in cognitive capacity, processing speed, and behavioral repertoire cooccur with changes in the structure and function of complex neural networks. Recent work has turned to the study of brain signal variability to inform our understanding of the processes underlying the formation of these complex neural networks. While inadequate or excessive neural variability provides inconsistent representations of the external world, which might result in poorly integrated neural networks and detrimental behavioral outcomes (Bosl et al., 2011, 2017; Catarino et al., 2011; Gurau et al., 2017; Sathyanarayana et al., 2020; Takahashi et al., 2010), a moderate amount of random noise in a system can, perhaps counterintuitively, enhance signal detection by improving the fidelity of an underlying signal (Figure 1) (Ward et al., 2006). Such variability is a fundamental property of neural systems at multiple hierarchical levels, and is thought to promote the exchange of information between neurons, neural synchrony, and the formation of robust, adaptable, and dynamic networks that are not overly reliant on any particular node (Fuchs et al., 2007; Miši et ć al., 2015; Shew et al., 2009; Ward et al., 2006). It is therefore increasingly understood that the inherently fluctuating nature of the brain, which is often modeled out of analyses as mere “noise,” serves a valuable functional role (Faisal et al., 2008; Garrett et al., 2013, 2011; Stein et al., 2005; Ward et al., 2006).

**Figure 1.**
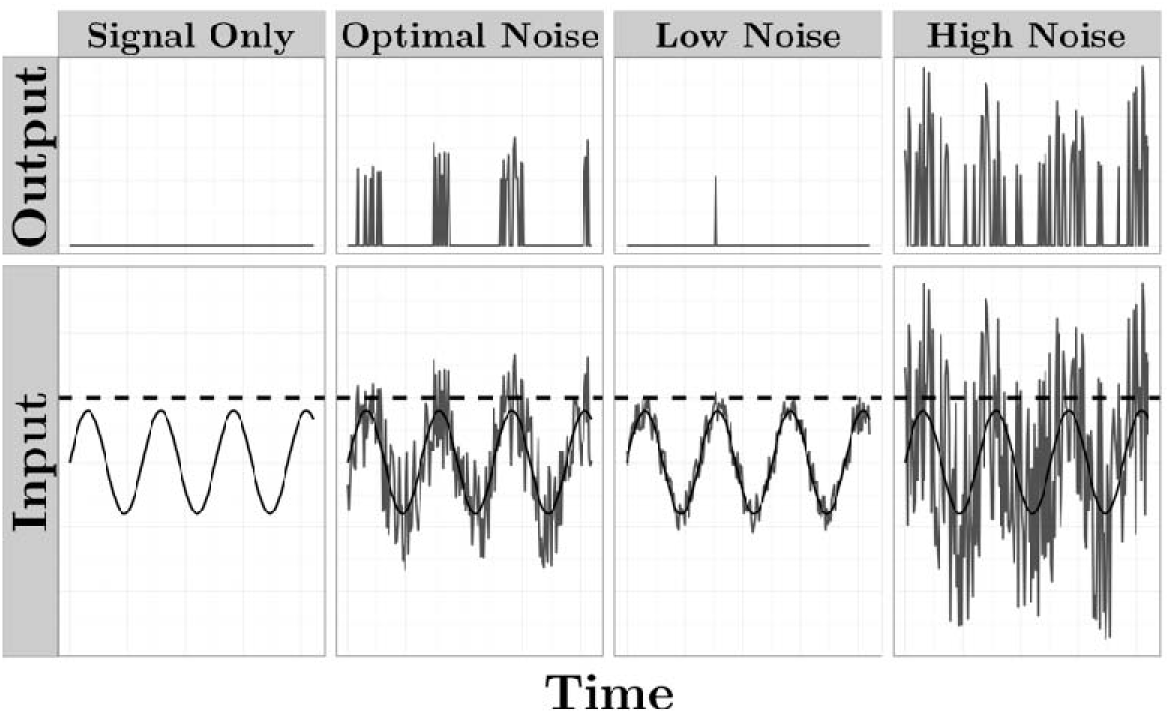
Adding random noise to a signal enhances signal detection. A theoretical illustration demonstrating that a signal that is below the threshold for detection (panel 1) can be enhanced and more accurately represented by the addition of a moderate amount of random noise (panel 2). However, inadequate (panel 3) or excessive (panel 4) noise provides inconsistent representations of the signal.

### Multiscale Entropy

The multiscale entropy (MSE) algorithm (Costa et al., 2005, 2002) is among the most popular methods to quantify such moment-to-moment brain signal variability by calculating entropy – a measure of irregularity or unpredictability – across multiple time scales. Entropy at fine time scales is understood to reflect local information processing, while entropy at coarser time scales relates to the long-range integration of information across distal neural nodes (Vakorin et al., 2013).

MSE computation involves 1) coarse graining the time series to scale *s* by averaging together *s* successive, non-overlapping data points, and 2) computing sample entropy (Richman and Moorman, 2000) on the resulting time series (Costa et al., 2002). Sample entropy quantifies irregularity by determining how frequently a pattern of length *m* repeats relative to a pattern of length *m*+1. A similarity criterion, *r*, set as a proportion of the standard deviation of the time series, determines what points are considered indistinguishable. For any data point *x*, all points within *x* ± *r* are considered indistinguishable for pattern matching. Then, the negative natural log of the ratio of the count of *m* patterns to the count of *m*+1 patterns is computed. Higher sample entropy values therefore indicate higher irregularity in the data because patterns of length *m*+1 reoccur less often than patterns of length *m* (Figure 2).

**Figure 2.**
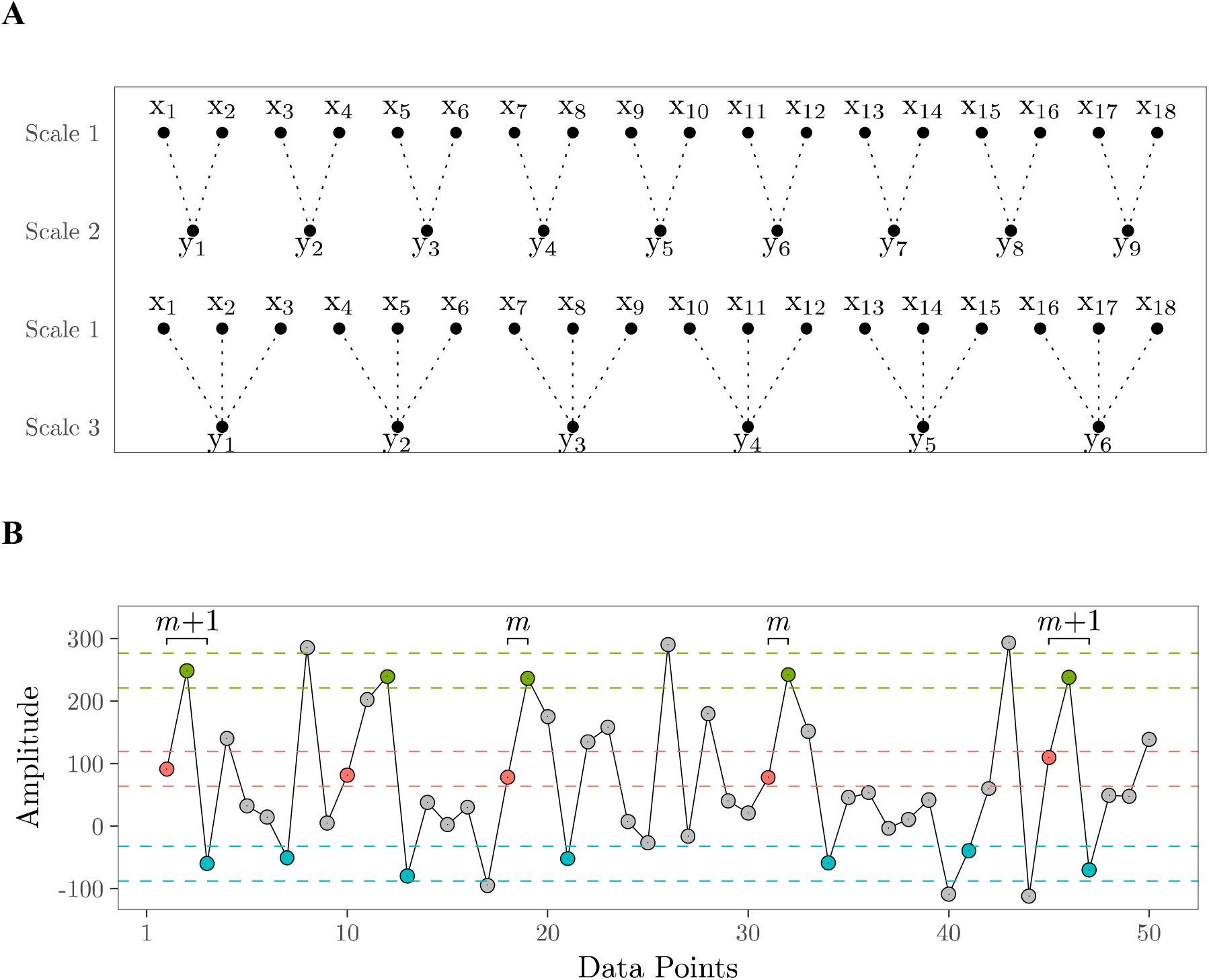
The multiscale entropy algorithm illustrated. (**A**) A coarse-grained time series is first computed for scale *s* by averaging together *s* consecutive, non-overlapping data points of the original time series (Scale 1). Entropy is then calculated on the coarse-grained time series. (**B**) Entropy measures the irregularity in a time series by determining how frequently a pattern of length *m* repeats relative to a pattern of length *m*+1. A similarity criterion, *r*, is set as a proportion of the standard deviation of the time series to determine which points are considered indistinguishable. For any data point *x*, all points within *x* ± *r* (illustrated with dashed lines) are considered indistinguishable. In this example, if *m* = 2, the first pattern of length *m* (points 1 and 2: red, green) repeats 4 times, whereas the first pattern of length *m*+1 (points 1, 2, 3: red, green, blue) repeats 2 times. The pattern template is then shifted forward 1 point such that matches of pattern *m* consisting of points 2 and 3, and pattern *m*+1 consisting of points 2, 3, and 4, are counted, and so on. Entropy is then calculated as the negative natural log of the ratio of the count of all pattern-length *m* repeats to the count of all pattern-length *m*+1 repeats: 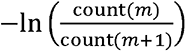 Consequently, low entropy values indicate regularity in a time series; if pattern length *m*+1 occurs as often as pattern length m, e.g.: 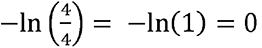 Conversely, high entropy values indicate high irregularity because patterns of length *m*+1 occur less often than patterns of length m, e.g.: 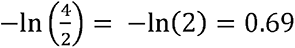.

In Costa’s original MSE algorithm (Costa et al., 2005, 2002), *r* is calculated as a percentage of the standard deviation of the original time series (i.e., scale 1) and remains constant across all scales. Using this method, it was shown that over increasingly coarse-grained time scales entropy increases for biological signals, such as heart rate or EEG data, but decreases for a completely random signal, such as white noise. It was argued that MSE was therefore capable of distinguishing truly “complex” time series from those that are completely random because “no new structures are revealed on larger scales” (Costa et al., 2005). However, a completely random time series should be highly irregular and unpredictable at any time scale, and therefore should yield high entropy values regardless of how the signal is coarse-grained. Instead, this decrease in entropy over time scales for random signals can be attributed to the fact that the standard deviation of a time series decreases with the coarse-graining procedure, and the extent of this decrease is greatest for random signals (Figure 3). Because sample entropy explicitly incorporates the standard deviation of the time series when defining the similarity criterion *r*, *r* is larger for a time series with greater standard deviation, meaning the entropy algorithm is more likely to identify matches resulting in a lower entropy value (Shafiei et al., 2019). Therefore, the original MSE algorithm conflates entropy with variance (Nikulin and Brismar, 2004). Recalculating the similarity criterion *r* at each time scale is a simple but critical modification to the MSE algorithm. Throughout the remainder of this article, we use “MSE” as an umbrella term to refer to any instance in which entropy is calculated across scales, but use “scale-wise entropy” to emphasize the importance of recalculating this parameter across scales and to differentiate when this modification is employed.

**Figure 3.**
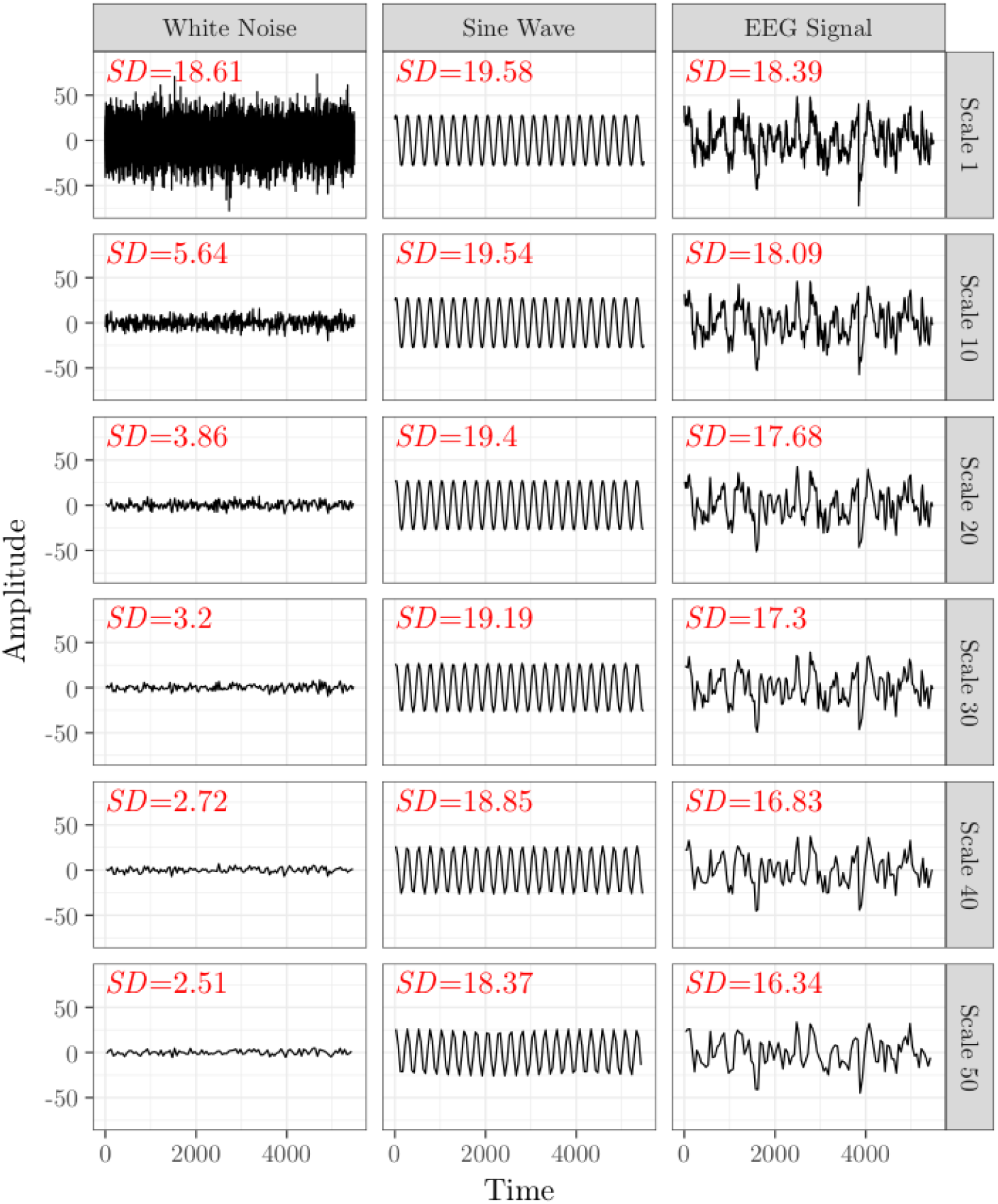
Coarse graining differentially impacts standard deviation across signal types. The original multiscale entropy curve involves setting the similarity criterion, *r*, as a proportion of the standard deviation (*SD*) of the native time series (Scale 1) and applying the parameter to all subsequent time scales. However, *SD* decreases as the scaling factor increases according to the statistical properties of the original time series. Here we plot a time series and its *SD* for simulated white noise (left), a sinusoidal wave (middle), and EEG signal (right) over scales 1, 10, 20, 30, 40, and 50. *SD* decreases most for white noise and least for the sine wave.

Calculating MSE from EEG signals requires careful consideration of data preprocessing procedures. EEG is susceptible to non-brain artifacts that themselves operate on different time scales, such as low frequency drifts and skin potentials, and high frequency muscle activity and electrical interference. Therefore, the typical preprocessing procedures applied to EEG data such as bandpass filtering to remove low and high frequency bands, and data cleaning procedures such as independent components analysis (ICA), require particular consideration for EEG data that will be subjected to MSE analysis. However, there is no standardized preprocessing protocol for the calculation of MSE on EEG data, and the preprocessing choices employed across different research labs vary widely (Table 1). For example, some argue that EEG data should undergo minimal preprocessing prior to MSE calculation to avoid the introduction of temporal distortions (e.g. (Okazaki et al., 2015)), while others maintain that non-brain sources of noise should be removed through thorough data cleaning procedures (e.g. (Miskovic et al., 2016)). Here, we develop and validate a standardized approach to preprocessing EEG data for the calculation of scale-wise entropy. We begin by reviewing previous work which has calculated MSE on pediatric EEG data to determine the range of preprocessing choices and the extent to which the critical modification to the MSE algorithm has been adopted (i.e., scale-wise entropy). We then select a representative range of preprocessing parameters and apply them to an infant EEG dataset to demonstrate the effect of preprocessing choices on scale-wise entropy estimates. Finally, we recommend a standardized approach to preprocessing and scale-wise entropy estimation that generates reliable scale-wise entropy estimates that are capable of differentiating developmental stages and cognitive states. Called the Automated Preprocessing Pipe-Line for the Estimation of Scale-wise Entropy from EEG Data (APPLESEED), this pipeline is made freely available as a fully automated and customizable MATLAB function that can be downloaded from https://github.com/mhpuglia/APPLESEED. The dataset used herein to develop and validate the pipeline is available for download from https://openneuro.org/datasets/ds003710 (Williams and Puglia, 2021).

**Table 1.** Summary of prior research applying MSE analysis to pediatric EEG data. The results of our literature review are summarized by key preprocessing and entropy algorithm parameters and the number of articles that employed each particular parameter including the sampling rate (native or after down-sampling), the method and frequency cutoffs for filtering, whether/which artifact correction methods were employed, the entropy *m*, pattern length parameter, the entropy *r*, similarity criterion parameter, and whether or not *r* was recalculated at each scale. Ns- parameter not specified. Numbers in brackets indicate the most frequently employed value.

| Article Count | Parameter |
| --- | --- |
| **Final sampling rate** |  |
| 4 | 1000 |
| 1 | 512 |
| 9 | 500 |
| 8 | 256 |
| 8 | 250 |
| 5 | 200 |
| 1 | 167 |
| 1 | 128 |
| 2 | 125 |
| 3 | multiple |
| 1 | ns |
| **Filtering** |  |
| 31 | Bandpass<br>High-pass: 0-1.5 [.5]<br>Low-pass: 20-120 [40;50] |
| 5 | Notch 50; 60 [60] |
| 2 | Lowpass 40; 64 |
| 5 | ns |
| **Artifact correction** |  |
| 7 | ICA + Interpolation |
| 3 | ICA |
| 33 | None/ns |
| ***m*** |  |
| 33 | 2 |
| 1 | 3 |
| 1 | 4 |
| 8 | ns |
| ***r*** |  |
| 9 | 0.15 |
| 17 | 0.20 |
| 2 | 0.25 |
| 6 | 0.50 |
| 9 | ns |
| **Scale-wise *r*** |  |
| 2 | Yes |
| 41 | No/ns |

### MSE Across Development: An Overview of Prior Pediatric EEG Research

To gain a comprehensive picture of the different preprocessing steps undertaken in the quantification of MSE in pediatric EEG, we conducted a literature search by entering the search terms (“multiscale entropy” OR “multi-scale entropy” OR “MSE” OR “multi scale entropy” OR “sample entropy”) AND (“EEG” OR “electroencephalography”) AND (“infan*” OR “newborn” OR “neonate” OR “child*” OR “adolescen*” OR “pediatric” OR “juvenile” OR “toddler” OR “developmental”) into PubMed and Web of Science databases. This search revealed 98 unique articles, 40 of which met inclusion criteria for our review. Articles were included if they were written in English and described original research in which multiscale entropy was estimated from EEG data collected in a pediatric (≤ 16 years of age) sample. Three additional articles identified through the reference lists of identified articles were also included. We extracted EEG recording, data preprocessing, and MSE algorithm parameters from each article (summarized in Table 1), which informed the selection of preprocessing methodology examined here.

While many studies demonstrate change in MSE across development (Bosl et al., 2011; De Wel et al., 2017; Hasegawa et al., 2018; Kang et al., 2019; Lippé et al., 2009; McIntosh et al., 2008; Miskovic et al., 2016; Polizzotto et al., 2016; Szostakiwskyj et al., 2017; Zhang et al., 2009) or with developmental disorder (Begum et al., 2017; Chenxi et al., 2016; Eroğlu et al., 2020; Kang et al., 2018; Liu et al., 2017; Okazaki et al., 2015; Rezaeezadeh et al., 2020; Simon et al., 2017; Wadhera and Kakkar, 2020; Weng et al., 2017), the vastly differing preprocessing choices and widespread failure to adopt the critical MSE algorithm modification of scale-wise recalculation of the similarity criterion makes it challenging to compare results across studies and realize the role of entropy in neurodevelopment.

## Automated Preprocessing Pipe-Line for the Estimation of Scale-wise Entropy from EEG Data (APPLESEED)

Here, we introduce APPLESEED, the Automated Preprocessing Pipe-Line for the Estimation of Scale-wise Entropy from EEG Data, and validate this novel pipeline for the analysis of EEG data collected in pediatric populations. We use the term scale-wise entropy to emphasize that this pipeline adopts the critical modification to the MSE algorithm that recalculates the similarity criterion parameter across scales.

APPLESEED is a fully automated and customizable MATLAB (The Math Works, Natick, MA) function that makes use of the freely available EEGLAB software (Delorme and Makeig, 2004) and associated plugins. APPLESEED was developed and validated using MATLAB 2017b and functions and scripts from EEGLAB v2021.1 (Delorme and Makeig, 2004) (download link: https://sccn.ucsd.edu/eeglab/download.php), ERPLAB v8.10 (Lopez-Calderon and Luck, 2014) (available as an EEGLAB plugin, via EEGLAB > File > Manage EEGLAB extensions), MADE Pipeline v1.0 (Debnath et al., 2020) (download link: https://github.com/ChildDevLab/MADE-EEG-preprocessing-pipeline), ADJUST (Mognon et al., 2011) (available as an EEGLAB plugin), and FASTER v1.0 (Nolan et al., 2010) (available as an EEGLAB plugin).

### Setting up to use APPLESEED

APPLESEED scripts can be downloaded from https://github.com/mhpuglia/APPLESEED, and include APPLESEED_setup(), which demonstrates how to prepare raw data for use in APPLESEED, APPLESEED(), which executes the preprocessing and entropy estimation pipeline on the prepared dataset, and APPLESEED_batch, which demonstrates how APPLESEED can be run as a batch across multiple subjects’ data. The dataset from this article is available for download from https://openneuro.org/datasets/ds003710 (Williams and Puglia, 2021). This provided dataset is organized according to the standardized Brain Imaging Data Structure (BIDS) format (Gorgolewski et al., 2016; Pernet et al., 2019), and we recommend that users follow this convention for naming and organizing their datasets for use with APPLESEED. In short, each subject’s EEG data is named as sub-<*identifier*>[_ses-<*identifier*>]_task-<*identifier*>[_acq-<*identifier*>][_run-<*identifier*>]_eeg<.*extension*> (terms in brackets are optional, if applicable). EEG file(s) are saved within an “eeg” sub-directory within a (session-level, if applicable, then) subject-level directory, housed within a study-wide parent directory (e.g., the path to the Brain Vision header file for the first recording of the provided dataset is: APPLESEED Example Dataset > sub-01 > ses-1 > eeg > sub-01_ses-1_task-appleseedexample_eeg.vhdr).

APPLESEED requires an EEGLAB dataset with an associated channel location structure as its input. If the data are not already in this format, the user can use the APPLESEED_setup() function to first import raw EEG data into EEGLAB. APPLESEED_setup() will automatically import raw data that is in a BIDS-approved file format (i.e., European Data Format, BrainVision Core Data Format, MATLAB toolbox EEGLAB, or Biosemi). If data are not in these formats, the user may edit Step 3 of APPLSEED_setup() or use the EEGLAB graphical user interface (GUI) to import the data using one of EEGLAB’s data import plugins that are available for many file format types (see https://eeglab.org/tutorials/04_Import/Importing_Continuous_and_Epoched_Data.html). Next, APPLESEED_setup() will assign channel locations (https://eeglab.org/tutorials/04_Import/Channel_Locations.html) using EEGLAB’s pop_chanedit() function. Alternatively, the user may specify the channel location file via the optional ‘chanfile’ input argument to APPLESEED(). We provide an example channel location file for the present dataset (in the “APPLESEED Example Dataset > code” directory). The data is then saved as an EEGLAB dataset within a subject-level (and, if applicable, session-level) directory within a “derivatives” folder housed in the parent directory (e.g., APPLESEED Example Dataset > derivatives > sub-01 > ses-1 > eeg > sub-01_ses-1_task- appleseedexample_eeg.set).

### Running APPLESEED

APPLESEED is executed as a function from the MATLAB command line. Mandatory input arguments for APPLESEED() include the file name for the EEGLAB dataset created during setup, the full path to the study directory, and, if a task-based analysis, the full path to the location of an ERPLAB bin file. A bin file defines unique event codes in the dataset and how they should be grouped within a task condition. We provide an example bin file for the present dataset (in the “APPLESEED Example Dataset > code” directory) and refer users to ERPLAB’s documentation for specifics on creating a bin file (https://github.com/lucklab/erplab/wiki/Assigning-Events-to-Bins-with-BINLISTER:-Tutorial). If event codes are found and no bin file is specified, a warning message will be displayed and the data will be treated as continuous, resting state data.

The user may also specify additional, optional arguments to customize preprocessing parameters. The default parameters for these inputs are based on the recommendations from this manuscript. Table 2 provides a description of all possible APPLESEED() input arguments and the default values that will be assigned if the argument is not specified at the command line.

**Table 2.**
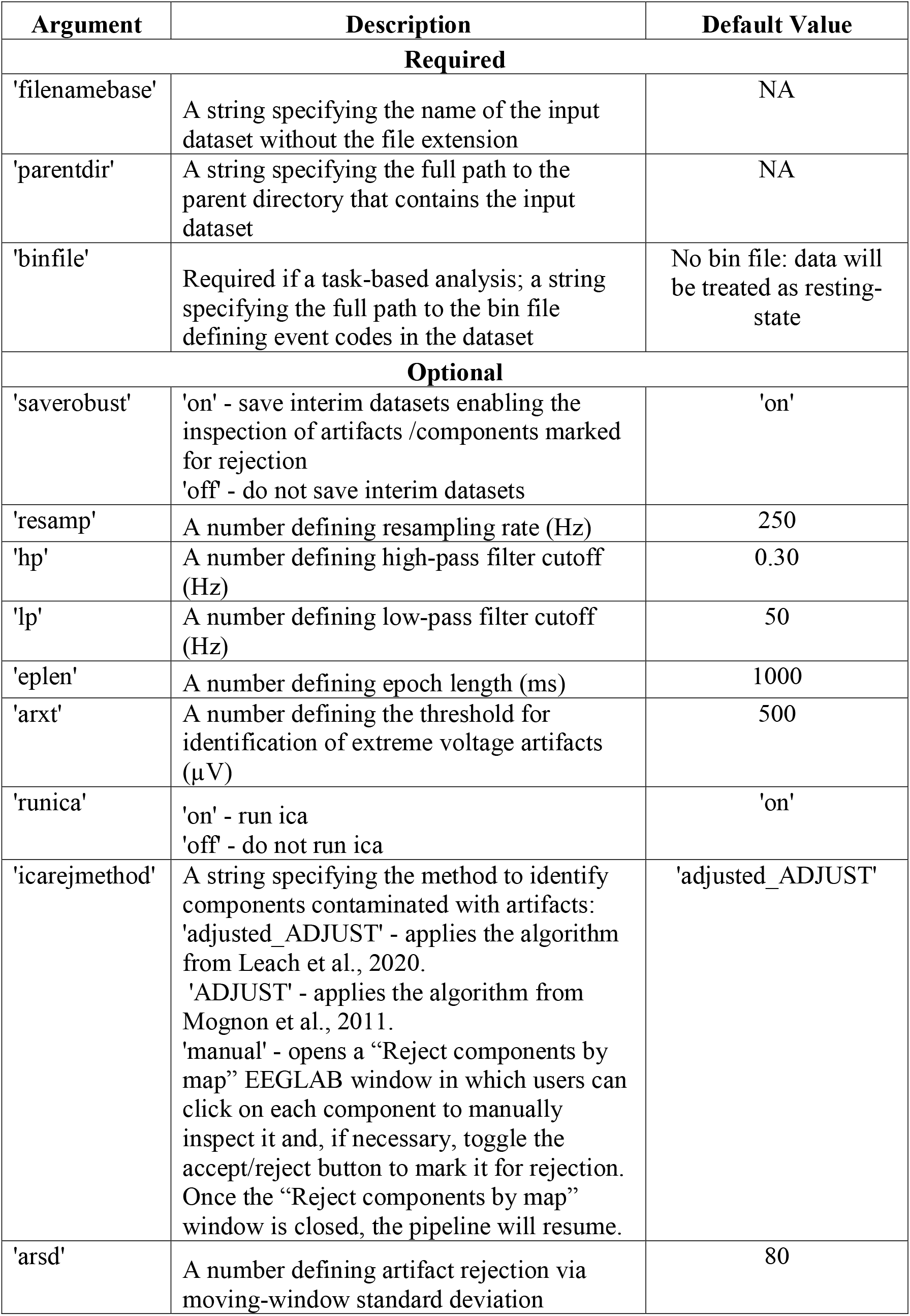
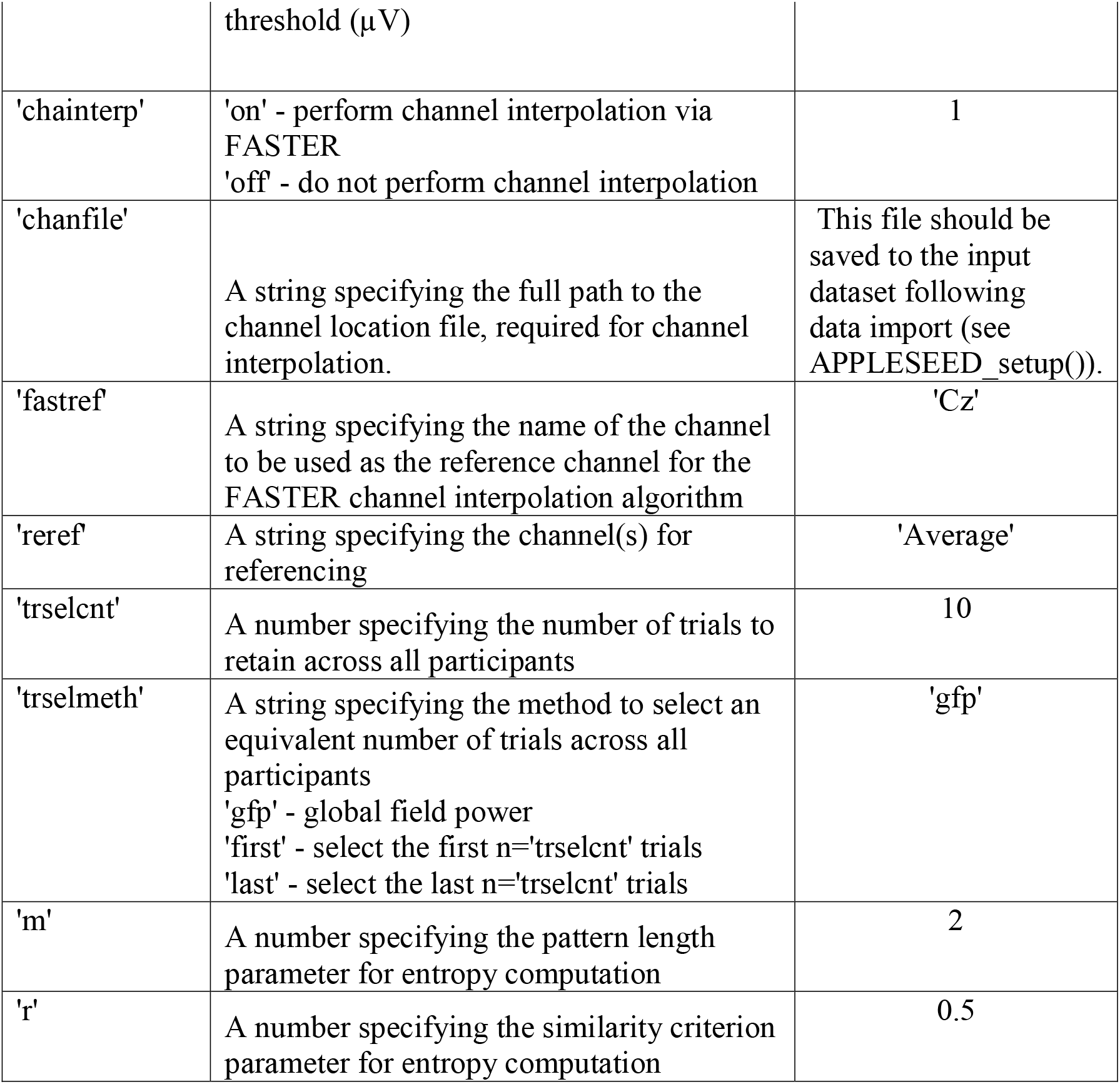
APPLESEED input arguments. . A description of all required and optional input arguments to the APPLESEED() function including the argument flag and the default value if the argument is not specified.

Output files are saved within a subject-level (and, if applicable, session-level) directory within an “appleseed” folder housed in the parent directory (e.g., APPLESEED Example Dataset > appleseed > sub-01 > ses-1 > eeg), and include the final, preprocessed dataset, a logfile detailing each preprocessing step employed and any errors or warnings that occurred during pipeline execution, scale-wise entropy file(s) (one per condition if a task-based analysis), and interim datasets that allow users to examine trials and components marked for rejection. While we strongly recommend that users inspect these datasets to ensure artifacts and components are appropriately classified, this option may be turned off via the optional ‘saverobust’ input argument.

The provided APPLESEED_batch script demonstrates how APPLESEED_setup() and APPLESEED() can be run as a batch across multiple subjects’ data, and runs APPLESEED for the present dataset (Williams and Puglia, 2021), which can be downloaded from https://openneuro.org/datasets/ds003710. Each step of APPLESEED is detailed below, and an overview of the pipeline is provided in Figure 4.

**Figure 4.**
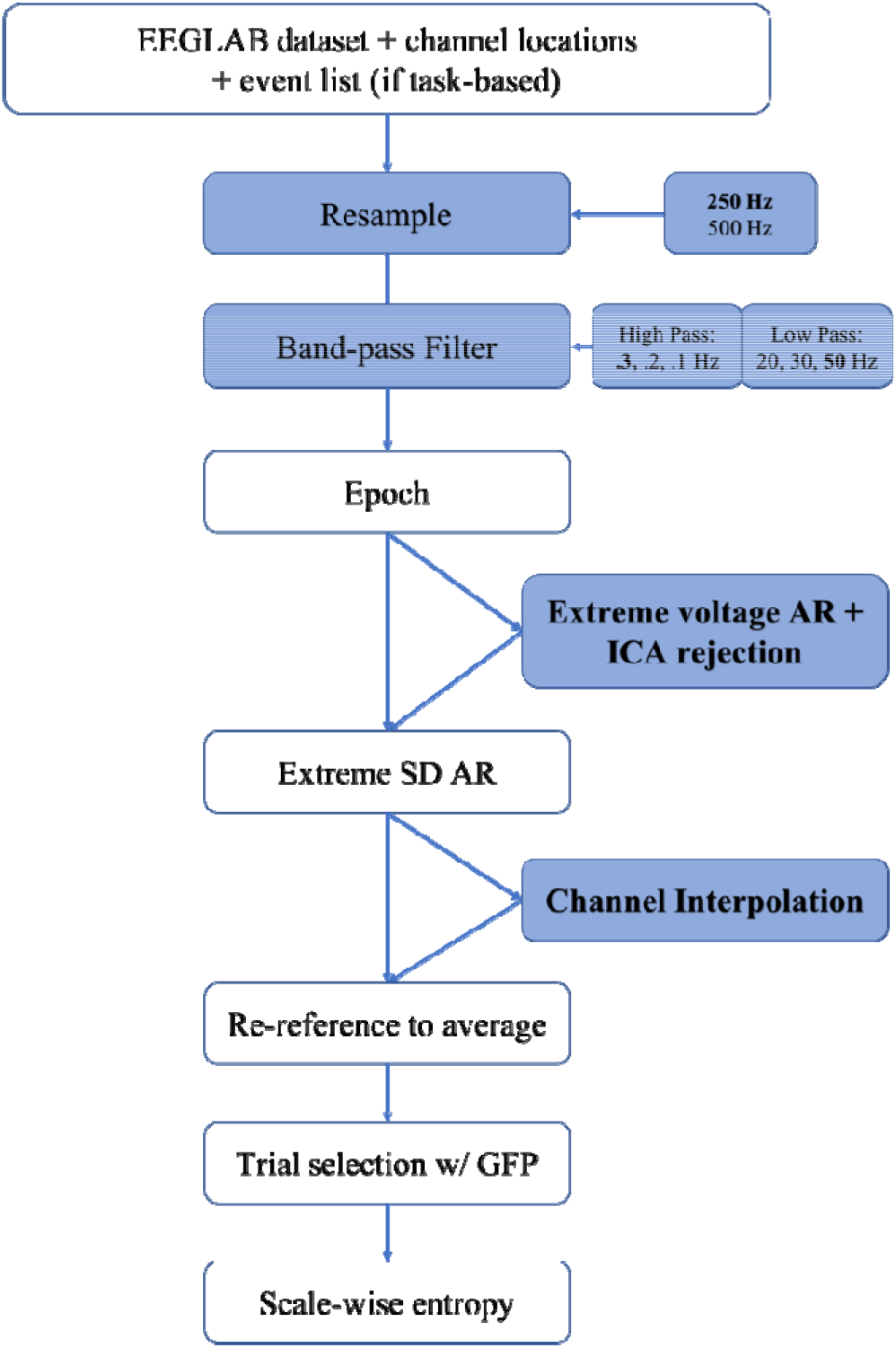
Overview of APPLESEED Preprocessing Pipeline. A flowchart depicting the preprocessing steps undertaken in APPLESEED. Blue coloring indicates steps for which alternative parameters were tested during the validation of this pipeline, and the boldfaced font indicates which parameter was ultimately selected for the optimized pipeline. AR – artifact rejection; SD – standard deviation; GFP – global field power.

### Preprocessing Step: Resampling

The first step in APPLESEED is to down-sample the data to a standardized sampling rate. In MSE, scales are directly related to the sampling rate of the native (scale 1) time series. For example, scale 1 for data sampled at 250 Hz, scale 2 for data sampled at 500 Hz, and scale 4 for data sampled at 1kHz comprise equivalent time scales. Therefore, scales cannot be directly compared across studies if different sampling rates are employed. For pipeline validation, we consider data down-sampled to 250 Hz and 500 Hz. Users may specify a resampling rate via the optional ‘resamp’ input argument.

### Preprocessing Step: Filtering

Filtering removes low-frequency drifts such as those associated with skin potentials, and high-frequency artifacts such as those introduced by muscle activity or electrical line noise. APPLESEED applies an infinite impulse response (IIR) Butterworth bandpass filter to the continuous EEG data. For pipeline validation, we consider high-pass cutoffs of 0.1, 0.2, and 0.3 Hz and low-pass cutoffs of 20, 30, and 50 Hz. Users may specify high- and low-pass cutoffs via the optional ‘hp’ and ‘lp’ input arguments, respectively.

### Preprocessing Step: Epoching

Next, the data is segmented into discrete epochs. For task-based studies, epochs are time-locked to stimulus onset. For resting-state studies, evenly-spaced epochs are extracted from the continuous time series. The default epoch length for APPLESEED is 1000 ms. We recommend using the longest possible epoch that enables an appropriate balance between artifact rejection and subject retention for subsequent analysis. While shorter epochs are less likely to contain eye blink or motion artifacts, epochs must be long enough to contain sufficient continuous data points for a reliable estimation of entropy (Grandy et al., 2016) and to achieve the desired coarse-grained scales. Because the coarse-graining procedure employs a moving window, the number of data points decreases as a function of scale. Users may specify an epoch length (in ms) via the optional ‘eplen’ input argument.

### Optional Preprocessing Step: ICA Rejection

Data may then be cleaned via artifact correction including ICA decomposition and channel interpolation. Because ICA performs best with large amounts of relatively clean data (Luck, 2014), epochs with extreme voltages (default threshold ± 500 µV, users may specify an alternate value via the ‘arxt’ input argument) are first rejected as artifacts. Data is then subjected to ICA decomposition using the EEGLAB runica() function. Components contaminated with artifacts must then be identified and removed. This identification may be performed manually (Lippé et al., 2009; McIntosh et al., 2008; Puglia et al., 2020) – a time intensive and somewhat subjective process, or via an automated algorithm. While several automated algorithms for the identification of artifactual components exist, APPLESEED makes use of the MADE (Debnath et al., 2020) adjusted_ADJUST function, which is the only algorithm specifically designed to detect artifactual components in pediatric data (Leach et al., 2020). This algorithm is an adaptation of the ADJUST EEGLAB plugin (Mognon et al., 2011), which examines the spatial and temporal features of each component to identify components contaminated by blinks, eye movements, and generic discontinuities. The MADE adjusted_ADJUST function makes several important modifications to improve performance of this algorithm on pediatric data including improved eye blink detection and retaining any components that contain an alpha peak (Debnath et al., 2020). For pipeline validation we consider both “maximal” data cleaning (i.e., ICA rejection + channel interpolation) and no data cleaning. Users may specify whether to run ICA and which algorithm (’adjusted_ADJUST’, ‘ADJUST’, or ‘manual’) to use for the automatic identification of components for rejection via the optional ‘runica’ and ‘icarejmethod’ input arguments, respectively.

### Preprocessing Step: Artifact rejection

Epochs with excessive amplitude standard deviations within a 200-ms sliding window with a 100-ms window step are discarded as artifacts. The default threshold value is set to 80_μV. If visual inspection reveals too many non-artifact epochs are rejected, users may wish to decrease this value, or if too many epochs with artifacts are retained, users may wish to increase this value. Users may specify a voltage threshold (in μV) for artifact rejection via the optional ‘arsd’ input argument.

### Optional Preprocessing Step: Channel Interpolation

Data may be further cleaned by channel interpolation. Problematic channels are identified and removed using the channel_properties() function from the FASTER EEGLAB plugin (Nolan et al., 2010). For each channel, this function computes and standardizes the channel’s correlation with other channels, the channel variance, and the channel’s Hurst exponent – a measure of long-range dependence within a signal (Nolan et al., 2010). If the value of one of these parameters exceeds 3 standard deviations from the mean, that channel is interpolated. APPLESEED’s default reference channel for the FASTER algorithm is Cz. Users may specify whether to run channel interpolation and an alternate FASTER reference channel via the optional ‘chaninterp’ and ‘fastref’ input arguments, respectively.

### Preprocessing Step: Re-referencing

Data are then re-referenced. By default, APPLESEED re-references to the average of all scalp electrodes, but users may specify an alternate re-referencing channel(s) via the optional ‘reref’ input argument.

### Preprocessing Step: Trial Selection

Finally, because the number of data points included in MSE calculation can influence the reliability of the estimates (Grandy et al., 2016), the final step of APPLESEED is trial selection of an equivalent number of trials across all participants by identifying the trials with total global field power (GFP) (McIntosh et al., 2008) closest to the median GFP for each participant. By default, APPLESEED will select 10 trials, but we recommend increasing this number as much as possible such that doing so retains a sufficient number of participants for subsequent analyses. Users may specify the number of trials to retain and the method for selecting these trials via the optional ‘trselcnt’ and ‘trselmeth’ input arguments, respectively.

### Scale-wise Entropy Calculation

Scale-wise entropy is then calculated using these selected trials for all electrodes in each dataset. To orthogonalize signal mean and signal variance, APPLESEED computes sample entropy on the residuals of the EEG signal (i.e., after subtracting the within-person average response across trials within each condition) using an algorithm based on that created by Grandy and colleagues for the estimation of MSE across discontinuous epochs (Grandy et al., 2016).

The default parameter values for entropy estimation are set to pattern length *m* = 2 and similarity criterion *r* = .5. Others have examined the effect of alternative *m* and *r* parameter values and found no substantial effect on the accuracy and precision of MSE estimates (Grandy et al., 2016). APPLESEED coarse-grains each time scale via moving window average, as in the original MSE algorithm. Critically, in scale-wise entropy, APPLESEED recalculates *r* for each scale. Users may specify alternative *m* and *r* values for entropy estimation via the optional ‘m’ and ‘r’ input arguments, respectively.

## Pipeline Development & Validation

To develop and validate APPLESEED, we iteratively applied multiple preprocessing parameters to an infant dataset (Williams and Puglia, 2021) that can inform both the test-retest reliability and the early developmental trajectory of scale-wise entropy. This dataset can be downloaded from https://openneuro.org/datasets/ds003710.

### Sample

As part of a larger, ongoing longitudinal study in which infants undergo EEG at 4, 8, and 12 months of age (Puglia et al., 2020), 14 infants were invited to return to the lab for EEG assessment within 1 week of their initial 4-month-old visit to establish the test-retest reliability of scale-wise entropy estimates. The primary caregiver accompanied the infant to all appointments and provided written informed consent for a protocol approved by the University of Virginia Health and Human Sciences Institutional Review Board. The target sample size for this test-retest reliability sample was determined via power analysis tables provided by Bujang and Baharum (Bujang and Baharum, 2017), which specify that 13 subjects are sufficient to detect an interclass correlation coefficient (*ICC*) of .70 based on two observations with 90% power. Retest data from one subject was of insufficient quality for analysis. Two participants failed to return for longitudinal assessment, and the data from one participant was of insufficient quality for analysis at subsequent visits. Therefore, the final dataset consists of 48 recording sessions, with reliability and longitudinal data for 11 infants (6 F), and reliability data, only, for an additional 2 infants (5 F). At the 4-month visit, infants ranged in age from 118 to 148 days (*M* = 129.14). The time between the test and retest appointments ranged from 1 to 8 days (*M* = 5.54). At retest, infants ranged in age from 124 to 155 days (*M* = 134.5). Infants ranged in age from 219 to 254 days (*M* = 241.18) at the 8-month visit, and from 334 to 427 days (*M* = 366.64) at the 12- month visit.

### EEG Acquisition

The present analyses make use of visual trials in which the infants viewed dynamic, colorful 2400-ms video clips of faces and objects in alternating 18-s blocks. Across conditions, stimuli were matched on low-level stimulus properties including luminance, contrast, spatial frequency, and visual angle (Puglia et al., 2020). EEG was recorded from 32 Ag/AgCl active actiCAP slim electrodes (Brain Products GmbH, Germany) affixed to an elastic cap according to the 10–20 electrode placement system. EEG was amplified with a BrainAmp DC Amplifier and recorded using BrainVision Recorder software with a sampling rate of 5L kHz. Infants were seated on their caregiver’s lap while undergoing EEG. Following a procedure widely used in developmental EEG experiments (Hoehl and Wahl, 2012), recording was terminated when the infant became fussy or inattentive. Participants successfully completed 4 to 12 blocks (*M* = 14.10) of each condition, and each block consisted of 6 stimuli.

Using this pipeline, we iteratively applied multiple preprocessing parameters to each infants’ dataset to determine what combination of parameters yielded reliable scale-wise entropy estimates that are sensitive to developmental changes and cognitive state. We applied the following preprocessing parameters to each infant’s EEG data: sampling rate (250 Hz, 500 Hz), high (0.1 Hz, 0.2 Hz, 0.3 Hz)- and low (20 Hz, 30 Hz, 50 Hz)-pass filter cutoffs, and whether artifact correction, i.e. via ICA and channel interpolation, was performed (no, yes), for a total of 36 preprocessing iterations prior to scale-wise entropy calculation.

To reduce the number of features considered, we averaged scale-wise entropy estimates across electrode regions of interest (ROIs, Figure 5) and frequency bands (see Table 3). The Frontal ROI consisted of electrodes Fp1, Fp2, F7, F3, Fz, F4, F8, FC5, FC1, FC2, and FC6. The centro-temporal ROI consisted of electrodes T7, C3, Cz, C4, T8, TP9, CP5, CP1, CP2, CP6, and TP10. The parieto-occipital ROI consisted of electrodes P7, P3, Pz, P4, P8, PO9, O1, Oz, O2, and PO10.

**Figure 5.**
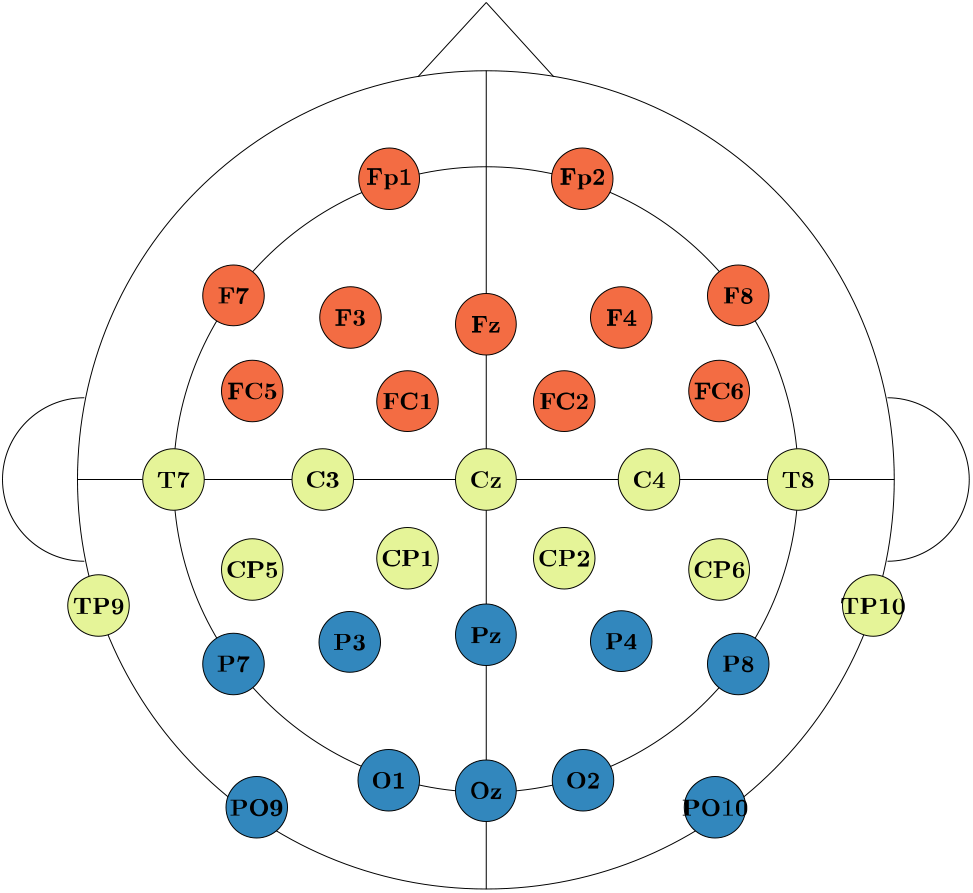
Electrode Cap Montage & Regions of Interest. EEG was recorded from 32 channels aligned according to the standard 10-20 system. To reduce the number of features considered, scale-wise entropy was averaged across electrode regions of interest (ROIs). These include the frontal ROI (red), the centro-temporal ROI (yellow), and the parieto-occipital ROI (blue).

**Table 3.** The summarization of scale-wise entropy estimates by frequency bands. For both considered sampling rates (250 Hz, 500 Hz), the scale range and total number of scales (n) that fell within each frequency band.

| Frequency Band | Hz | 250 Hz Scales (n) | 500 Hz Scales (n) |
| --- | --- | --- | --- |
| Delta | < 4 | 63 - 83 (21) | 126 - 166 (41) |
| Theta | 4 - 7 | 32 - 62 (31) | 63 - 125 (63) |
| Alpha | 8 - 12 | 20 - 31 (12) | 39 - 62 (24) |
| Beta | 13 - 29 | 9 - 19 (11) | 17 - 38 (22) |
| Gamma | 30 - 100 | 3 - 8 (6) | 5 - 16 (12) |
| Gamma+ | 101+ | 1 - 2 (2) | 1 - 4 (4) |

### Effect of Preprocessing on Data Retention

The accuracy and precision of sale-wise entropy estimates increases as a function of the number of data points included in the calculation (Grandy et al., 2016). Furthermore, longer time series enable the investigation of coarser time scales reflective of long-range integration (Vakorin et al., 2013). However, particularly within pediatric samples, EEG recordings are likely to be of short duration and contaminated with motion artifacts, yielding fewer usable trials. We therefore first examine how the proportion of data retained after preprocessing varies as a function of preprocessing parameters. Across all 36 preprocessing pipelines considered, the number of retained epochs after preprocessing ranged from 15 to 64 (*M* = 35.29) for the Viewing Faces condition, and from 10 to 58 (*M* = 35.03) for the Viewing Objects condition.

We entered proportion of data retained after preprocessing into a repeated measures analysis of variance (ANOVA) using the aov function within R (R Core Team, 2020), with sampling rate (250 Hz; 500 Hz), high-pass filter cutoff frequency (0.1 Hz, 0.2 Hz, 0.3 Hz), low-pass filter cutoff frequency (20 Hz, 30 Hz, 50 Hz), data cleaning implementation (no, yes), experimental condition (viewing faces, viewing objects), and study visit (1, 2, 3) as within-subjects factors. Proportion of data retained differed significantly by sampling rate (F(1,10) = 7.01, p = 0.024) such that more data was retained for data sampled at 500 Hz, high-pass (F(2,20) = 39.54, p < 0.001) and low-pass filter cutoffs (F(2,20) = 19.58, p < 0.001) such that more aggressive filters were associated with a greater proportion of the data retained, and data cleaning implementation (F(1,10) = 93.39, p < 0.001), such that more data was retained when data cleaning procedures were implemented (Figure 6). Proportion of data retained did not differ significantly by experimental condition (F(1,10) = 0.03, p = 0.872) or across longitudinal visits (F(3,30) = 1.24, p = 0.311). Average MSE curves for each preprocessing pipeline can be viewed in Figure 7. High-pass filter cutoff impacts mid-range time scales such that a higher cutoff filter is associated with higher entropy values, regardless of sampling rate or data cleaning implementation (Figure 7A). Low-pass filter cutoff impacts fine-grained time scales such that a higher cutoff filter is associated with higher entropy values, with a steeper initial slope for the lower sampling rate, regardless of data cleaning implementation (Figure 7B).

**Figure 6.**
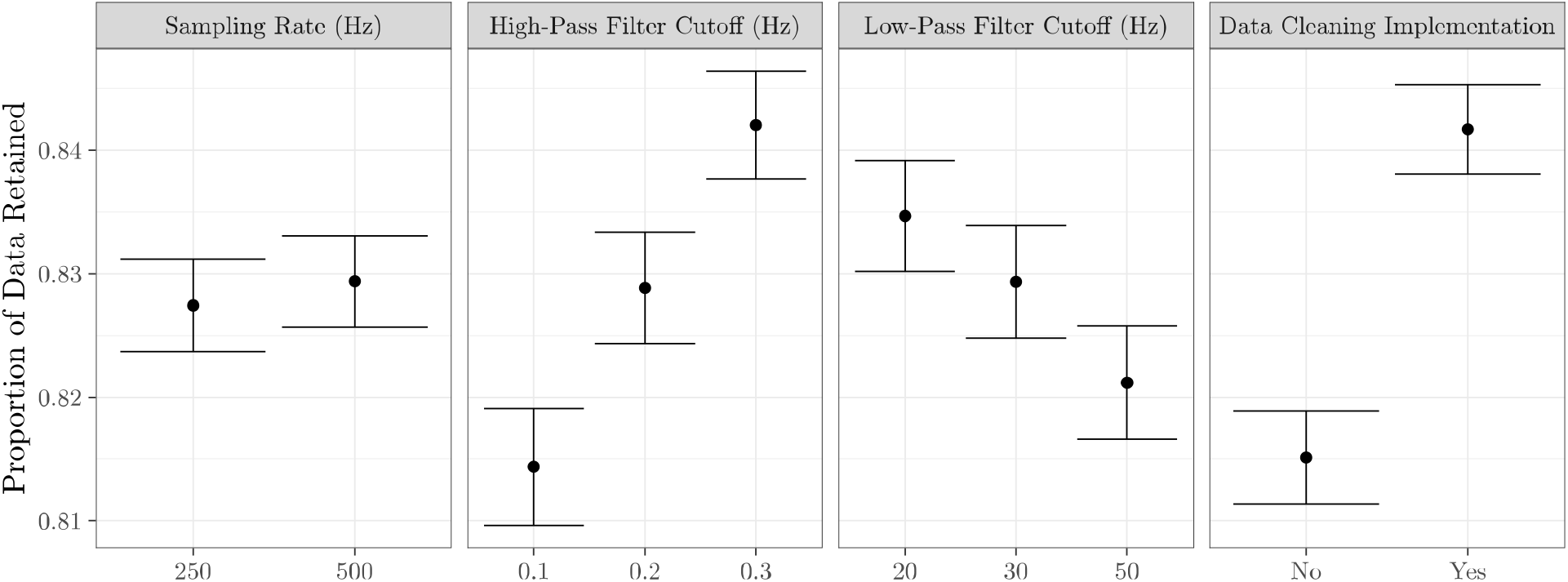
Proportion of data retained across preprocessing parameters. Results from a repeated measures ANOVA revealed that proportion of data retained significantly varies by sampling rate, high- and low-pass filter cutoffs, and data cleaning procedures.

**Figure 7.**
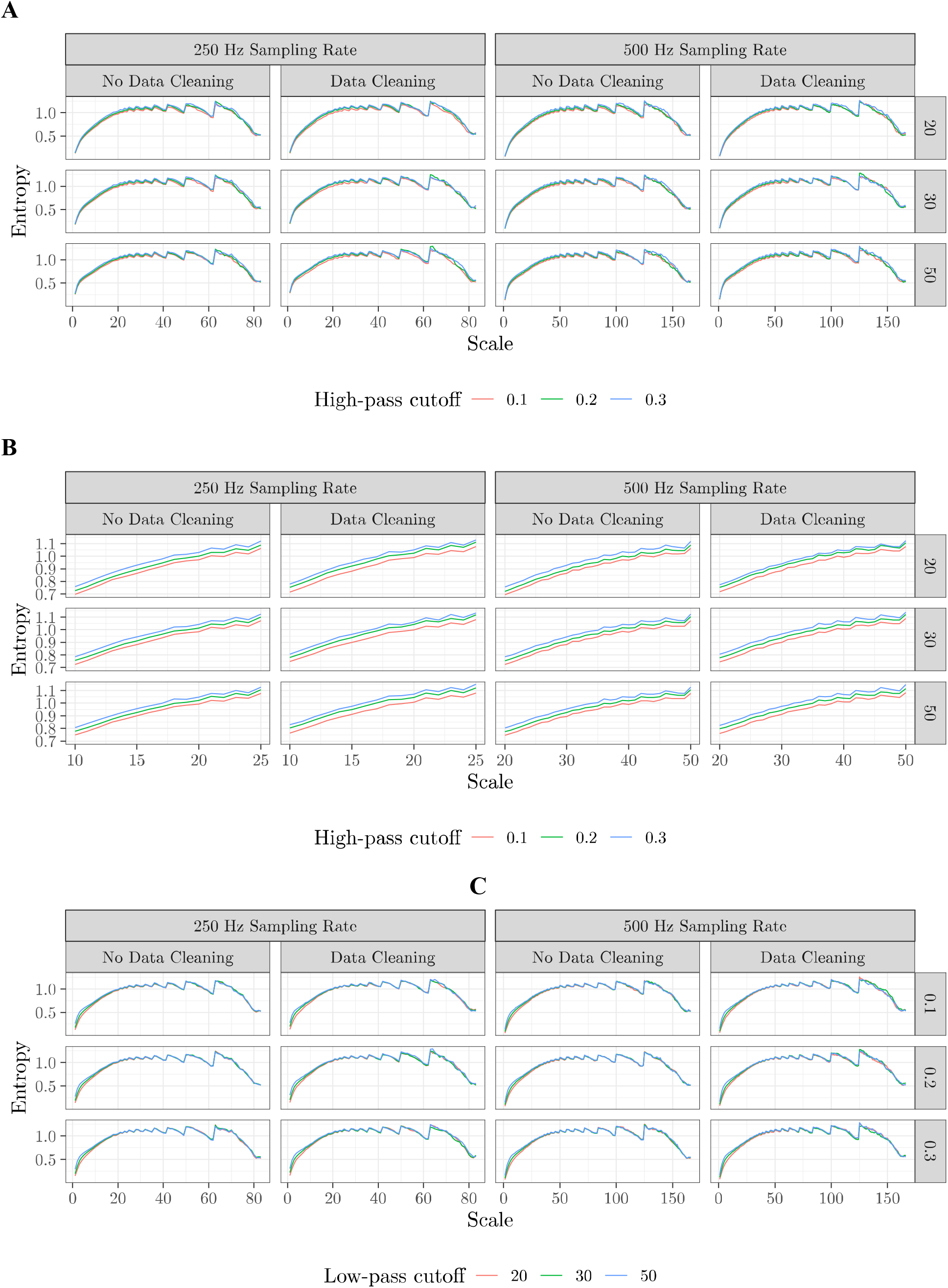
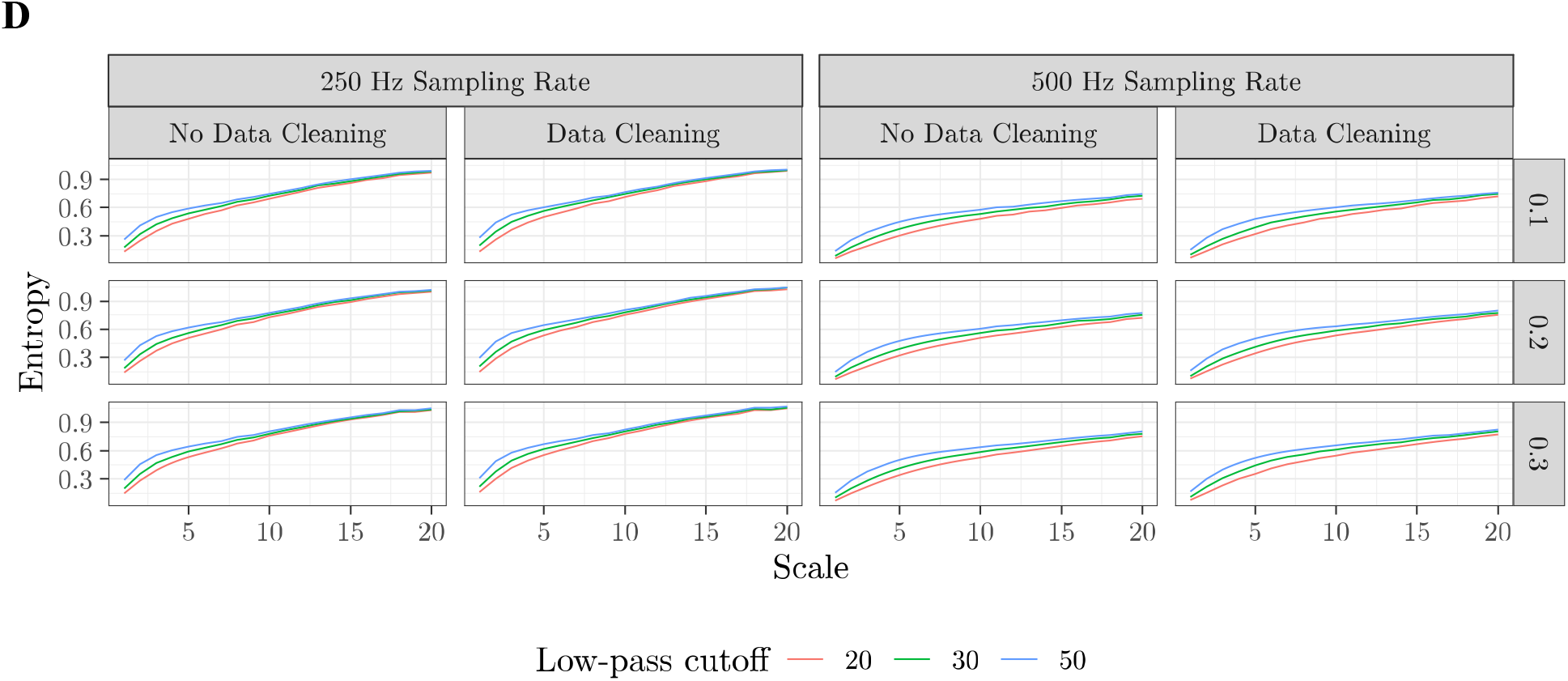
Impact of preprocessing procedures on scale-wise entropy curves. Scale- wise entropy is plotted as a function of sampling rate and data cleaning implementation for (**A,B**) High-pass filter cutoffs and (**C,D**) low-pass filter cutoffs considered in the development and validation of our pipeline. **A** & **C** depict scale-wise entropy curves for all considered scales. **B** & **D** depict a zoomed-in view for specific scales that depict the maximum impact of filtering.

### Reliability of Scale-wise Entropy Estimates

To develop and validate a standardized methodology for preprocessing pediatric EEG data for scale-wise entropy analysis, we first determine the reliability of scale-wise entropy estimates following different preprocessing procedures. We first calculate ICC using the icc function of the irr R package (Gamer et al., 2019) on overall scale-wise entropy estimates averaged across scales and ROIs for each condition. Only reliable estimates that are significantly reproducible in both experimental conditions are considered. Eight preprocessing pipelines yielded reliable ICC estimates across both Viewing Faces and Viewing Objects conditions (Table 4). ICC estimates for all bands, electrodes, and preprocessing parameters can be seen in Figure 8. Test-retest reliability curves for the final, recommended pipeline can be viewed in Figure 9A.

**Figure 8.**
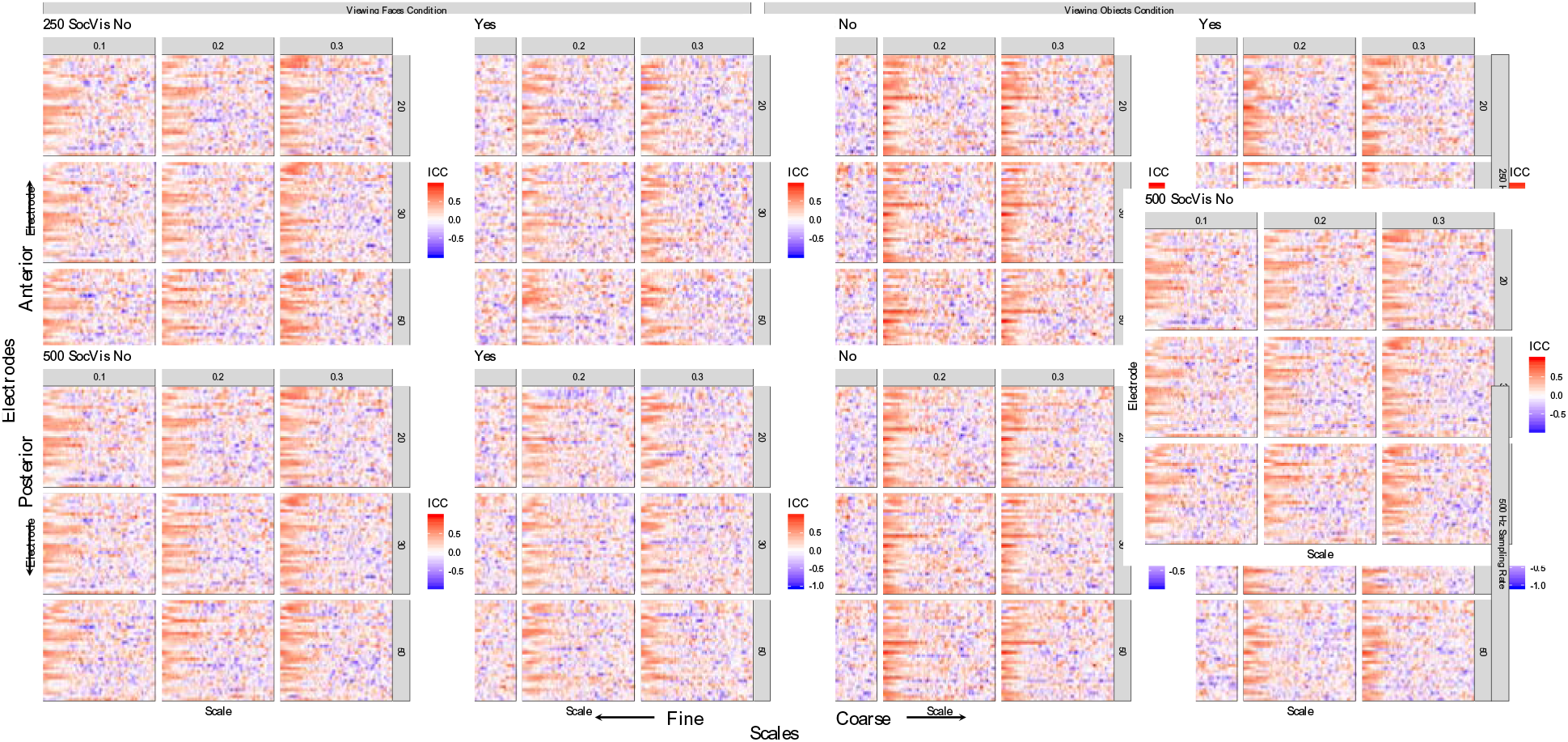
Test-retest reliability estimates across preprocessing parameters. The intraclass correlation coefficient (*ICC*) assessing the reliability of scale-wise entropy from the 4-month visit to the retest visit (approximately 1 week later) is plotted for each scale, electrode, and preprocessing parameter. In general, finer scales have higher reliability estimates across electrodes. Hotter colors represent higher *ICC*s.

**Figure 9.**
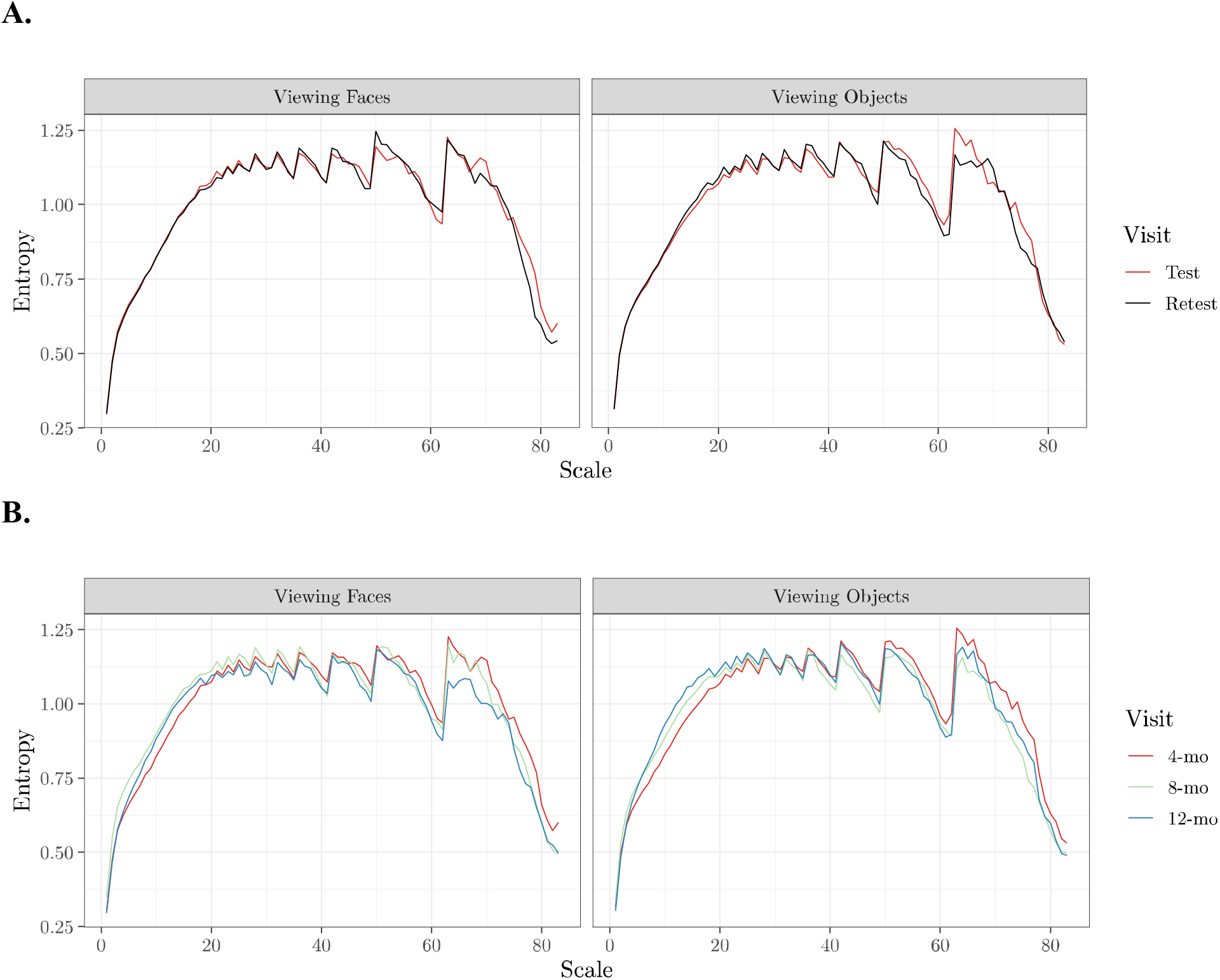
Scale-wise entropy curves generated with APPLESEED. **A.** Average test-retest reliability scale-wise entropy curves for each condition generated with the final preprocessing pipeline. **B.** Scale-wise entropy curves depicting the average developmental trajectory for each condition from 4- to 12-months of age.

**Table 4.** Preprocessing pipelines that produce reliable scale-wise entropy estimates. The eight preprocessing pipelines that yielded significantly reliable results across the test (4-months- of-age) and retest (within 1 week) visits in both the face viewing and object viewing conditions across all scales and all electrode ROIs. The values for the final, recommended APPLESEED parameters are highlighted in bold font. *ICC* – intraclass correlation coefficient; *p* – *p*-value.

| Sampling Rate (Hz) | High-Pass Cutoff (Hz) | Low-Pass Cutoff (Hz) | Artifact Correction | Viewing Faces Condition |  | Viewing Objects Condition |  |
| --- | --- | --- | --- | --- | --- | --- | --- |
|  |  |  |  | *ICC* | *p* | *ICC* | *p* |
| 250 | 0.2 | 50 | No | 0.49 | 0.041 | 0.55 | 0.016 |
| 250 | 0.3 | 20 | No | 0.45 | 0.039 | 0.45 | 0.047 |
| 250 | 0.3 | 20 | Yes | 0.53 | 0.019 | 0.51 | 0.034 |
| **250** | **0.3** | **50** | **Yes** | **0.47** | **0.048** | **0.55** | **0.024** |
| 500 | 0.2 | 50 | No | 0.46 | 0.047 | 0.59 | 0.010 |
| 500 | 0.3 | 20 | No | 0.41 | 0.048 | 0.50 | 0.027 |
| 500 | 0.3 | 30 | No | 0.46 | 0.035 | 0.48 | 0.037 |
| 500 | 0.3 | 50 | No | 0.63 | 0.007 | 0.44 | 0.050 |

### Scale-wise entropy Estimates are sensitive to developmental stage and cognitive state

Next, we examined how scale-wise entropy estimates change across development, and whether these estimates were capable of differentiating perceptual states across the two viewing conditions. For each frequency band and ROI, scale-wise entropy estimates were entered into repeated measures ANOVAs with within-subject factors of visit (1, 2, 3) and experimental condition (viewing faces, viewing objects). Greenhouse-Geisser correction was applied to any factors violating the assumption of sphericity (Mauchly’s test *p*-value ≤ 0.05). Preprocessing procedures that consistently yielded significant effects within at least 5 of the 6 frequency bands were considered further. Of these, one preprocessing pipeline overlapped with a preprocessing pipeline that generated reliable scale-wise estimates across conditions. When considering all electrodes, we find a significant main effect of age on scale-wise entropy estimates within the gamma+ (F(2,20) = 3.62, p = .045), gamma (F(2,20) = 3.61, p = .046), and delta (F(2,20) = 5.01, p = .017) frequency bands. In general, entropy increases from 4- to 8-months for fine-grained scales, but decreases within the delta frequency band over this time period (Figure 9A). We find a significant interaction between age and condition within the beta (F(2,20) = 4.37, p = .027) and alpha (F(2,20) = 3.71, p = .043) frequency bands. This interaction shows that there is no distinction between entropy estimates at the 4- and 8-month-old visits, but by the 12-month visit, entropy is capable of distinguishing between viewing conditions (Figure 9A).

When considering scale-wise entropy across frequency bands and ROIs, we find a significant main effect of age in frontal beta (F(1.33,13.26) = 6.08, p = .021), centro-parietal gamma+ (F(2,20) = 4.68, p= .021), gamma (F(1.27,12.7) = 9.93, p = .005), beta (F(1.19,11.94) = 9.66, p = .007), and delta (F(2,20) = 5.34, p = .014), and parieto-occipital theta (F(2,20) = 4.71, p = .021). Again, entropy generally increases from 4- to 8-months for fine-grained scales, but decreases over this time period for coarse-grained scales (Figure 10B). We also find a significant interaction between age and condition within parieto-occipital gamma (F(2,20) = 5.20, p = 0.015) and beta (F(2,20) = 4.75, p = 0.021). This interaction shows again that there is no distinction between entropy estimates at the 4- and 8-month-old visits, but by the 12-month visit, entropy within parietal and occipital regions, specifically, is capable of distinguishing between viewing conditions (Figure 10B). The developmental trajectory of scale-wise entropy as calculated by the final, recommended pipeline can be viewed in Figure 9B.

**Figure 10.**
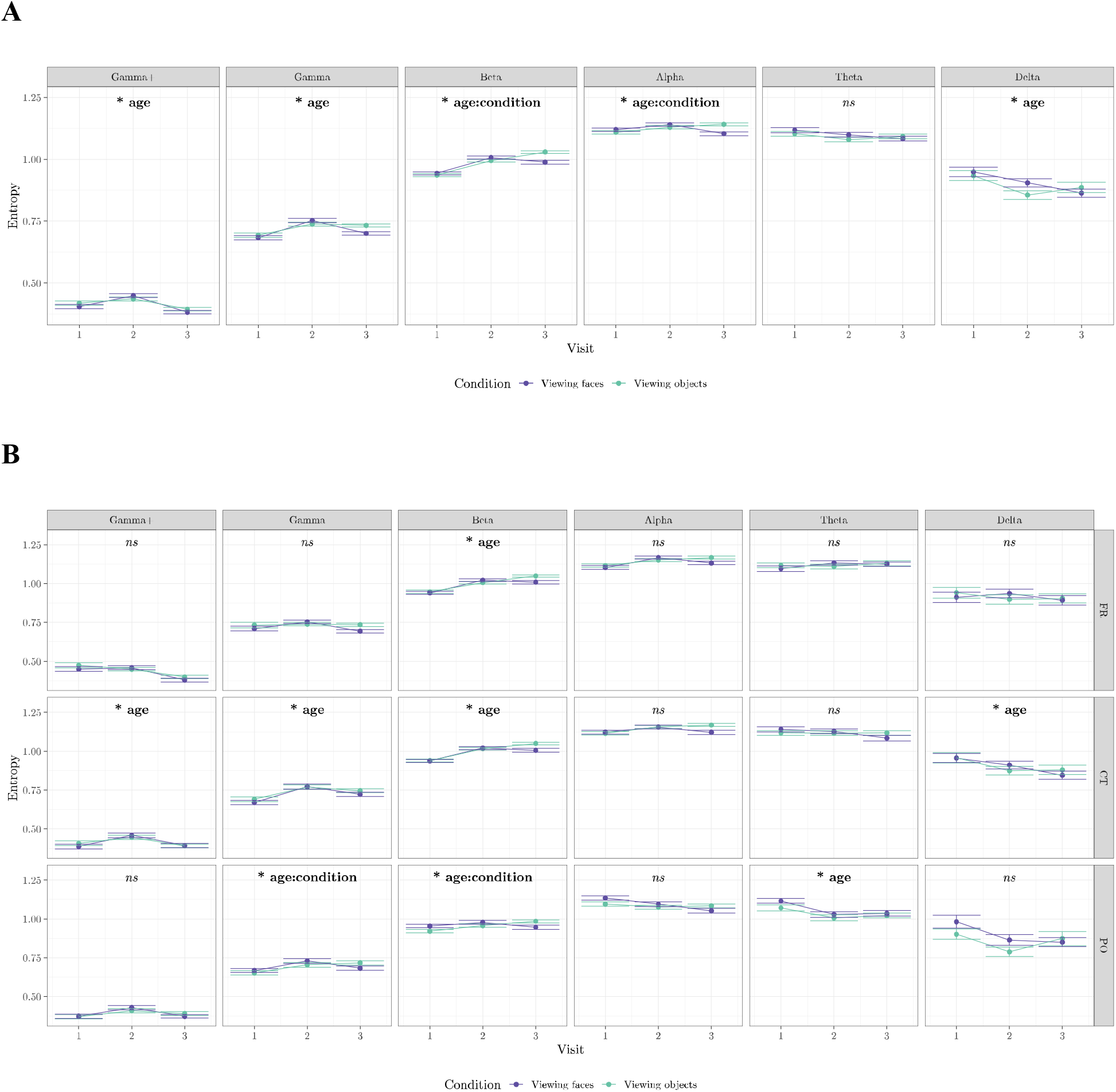
Scale-wise entropy across conditions and development. Results from a repeated measures ANOVA depicting the effect of age and condition on scale-wise entropy estimates across the whole brain (**A**) and within regions of interest (**B**) for each frequency band. Significant effects are indicated in each panel. ns – not significant; FR – frontal ROI; CT – centro-temporal ROI; PO – parieto-occipital ROI.

## Recommendations and Conclusions

We find a single preprocessing pipeline generates scale-wise entropy estimates that are both (1) significantly reliable across recording sessions occurring approximately 1 week apart in two experimental conditions, and (2) capable of differentiating cognitive states and developmental stages from 4- to 8-months-of-age. We therefore developed APPLESEED to automatically accomplish the following recommended preprocessing steps and scale-wise entropy estimation: data (down)sampling at 250 Hz, bandpass filtering with 0.3-50 Hz cutoffs, segmenting the data into 1000 ms (or longer, if possible) epochs, extreme artifact rejection, data cleaning via extreme artifact rejection, rejection of ICA components contaminated with artifacts via the automated adjusted_ADJUST algorithm, artifact rejection using a peak-to-peak moving window, channel interpolation of problematic channels identified via the FASTER package, re-referencing to the average of all scalp electrodes, and the selection of 10 (or more, if possible) trials across all participants via global field power. Finally, scale-wise entropy is calculated on the residuals of the EEG signal with pattern length *m* = 2 and similarity criterion *r* = 0.5 recalculated for each coarse-grained time scale.

While some prior work has examined the reliability and psychometric properties of MSE (Grandy et al., 2016; Kaur et al., 2019; Kuntzelman et al., 2018), these efforts employed the original, unmodified MSE algorithm that fails to recalculate *r* at each coarse-grained time scale – thereby conflating time series variance with entropy and hindering the ability to attribute any results to time series irregularity, specifically. We are the first to our knowledge to systematically examine the effect of preprocessing procedures, to make recommendations specifically for the use of scale-wise entropy in pediatric EEG datasets, and to provide freely available scripts to accomplish a standardized preprocessing pipeline for scale-wise entropy calculation adopting the critical variance-normalization algorithm modification.

### Limitations and future directions

The sample size for the present study was based on a power analysis for ICC estimation, and it cannot be overlooked that the sample size for the present study is small. While we may therefore be underpowered to detect condition-specific effects across developmental stages, it should be noted that our exploratory results align with hypothesized effects. Specifically, scale-wise entropy differentiates visual conditions in the parieto-occipital ROI beginning at 12-months of age. This result, in particular, highlights the plausibility of our results. Visual processing occurs in the occipital and parietal cortices, and we have previously shown scale-wise entropy associations within the visual domain do not yet emerge at 4 months of age in a larger sample (Puglia et al., 2020). These data suggest that brain signal entropy may be sensitive to developmental trajectories that aligns with sensory system maturation. Converging lines of research suggest that infants do not initially rely on visual cues for perception (Fernald, 1992; Mumme et al., 1996; Walker-Andrews, 1997). As with many mammals, the visual system matures later in development (Gottlieb, 1971), and in humans visual acuity does not reach adult levels until age 3 (Catford and Oliver, 1973).

Additionally, while APPLESEED is capable of processing resting state data, we only considered a task-based dataset consisting of passive viewing of visual stimuli for this initial validation study. Finally, we only considered the effects of sampling rate, high- and low-pass filter cutoffs, and a limited number of data cleaning algorithms. Alternative preprocessing procedures and entropy computation parameters may differentially impact results. For example, other coarse-graining methods may reveal alternative, complementary signatures of neural dynamics to the traditional moving-average window coarse-graining procedure employed here (Kosciessa et al., 2020). To overcome these limitations of the present study, and the limitations in interpreting prior results generated across a wide range of preprocessing procedures, we make APPLESEED freely available as a fully automated and customizable pipeline to facilitate future large-scale, multi-site investigations of scale-wise entropy effects throughout development using standardized, reproducible, and justified methods.

## Acknowledgments

The data supporting this manuscript are part of a larger study which was funded by N.S.F. grant 1729289 to Tobias Grossman, Jessica J. Connelly, and James P. Morris, and funds from NICHD grant F31HD090865, the American Psychological Foundation, the University of Virginia Brain Institute, and a PEO Scholar Award to M.H.P. We thank Tobias Grossmann for providing the facilities and equipment to collect the infant EEG data, Tobias Grossmann, Jessica J. Connelly, and James P. Morris for their contribution to the design of the original study, Kevin A. Pelphrey for providing feedback on this manuscript, Laura Atalla, Rachel Corney, Christiana King, Aaria Malhotra, Hannah Sharpe, and Rebecca Stafford for their assistance with data collection, and the families and infants who participated in this study.

## Bibliography

Begum, D., Ravikumar, K.M., Vykuntaraju, K.N., 2017. An initiative to classify different neurological disorder in children using multichannel EEG signals, in: 2016 IEEE International Conference on Recent Trends in Electronics, Information and Communication Technology, RTEICT 2016 - Proceedings. Institute of Electrical and Electronics Engineers Inc., pp. 1563–1566. https://doi.org/10.1109/RTEICT.2016.7808095

Bosl, W., Tierney, A., Tager-Flusberg, H., Nelson, C., 2011. EEG complexity as a biomarker for autism spectrum disorder risk. BMC Med. 9, 18. https://doi.org/10.1186/1741-7015-9-18

Bosl, W.J., Loddenkemper, T., Nelson, C.A., 2017. Nonlinear EEG biomarker profiles for autism and absence epilepsy. Neuropsychiatr. Electrophysiol. 3, 1. https://doi.org/10.1186/s40810-017-0023-x

Bujang, M.A., Baharum, N., 2017. A simplified guide to determination of sample size requirements for estimating the value of intraclass correlation coefficient: a review. Arch. Orofac. Sci. 12, 1–11.

Catarino, A., Churches, O., Baron-Cohen, S., Andrade, A., Ring, H., 2011. Atypical EEG complexity in autism spectrum conditions: A multiscale entropy analysis. Clin. Neurophysiol. 122, 2375–2383. https://doi.org/10.1016/j.clinph.2011.05.004

Catford, G. V, Oliver, A., 1973. Development of visual acuity. Arch. Dis. Child. 48, 47–50. https://doi.org/10.1136/adc.48.1.47

Chenxi, L., Chen, Y., Li, Y., Wang, J., Liu, T., 2016. Complexity analysis of brain activity in attention-deficit/hyperactivity disorder: A multiscale entropy analysis. Brain Res. Bull. 124, 12–20. https://doi.org/10.1016/j.brainresbull.2016.03.007

Costa, M., Goldberger, A.L., Peng, C.-K., 2002. Multiscale Entropy Analysis of Complex Physiologic Time Series. Phys. Rev. Lett. 89, 068102. https://doi.org/10.1103/PhysRevLett.89.068102

Costa, M., Goldberger, A.L., Peng, C.K., 2005. Multiscale entropy analysis of biological signals. Phys. Rev. E 71, 021906. https://doi.org/10.1103/PhysRevE.71.021906

De Wel, O., Lavanga, M., Caicedo Dorado, A., Jansen, K., Dereymaeker, A., Naulaer, G., Van Huffel, S., 2017. Complexity Analysis of Neonatal EEG Using Multiscale Entropy: Applications in Brain Maturation and Sleep Stage Classification. Entropy 19.

Debnath, R., Buzzell, G.A., Morales, S., Bowers, M.E., Leach, S.C., Fox, N.A., 2020. The Maryland analysis of developmental EEG (MADE) pipeline. Psychophysiology 57, e13580. https://doi.org/10.1111/psyp.13580

Delorme, A., Makeig, S., 2004. EEGLAB: An open source toolbox for analysis of single-trial EEG dynamics including independent component analysis. J. Neurosci. Methods 134, 9–21. https://doi.org/10.1016/j.jneumeth.2003.10.009

Eroğlu, G., Gürkan, M., Teber, S., Ertürk, K., Kırmızı, M., Ekici, B., Arman, F., Balcisoy, S., Özgüz, V., Çetin, M., 2020. Changes in EEG complexity with neurofeedback and multi- sensory learning in children with dyslexia: A multiscale entropy analysis. Appl. Neuropsychol. Child. https://doi.org/10.1080/21622965.2020.1772794

Faisal, A.A., Selen, L.P.J., Wolpert, D.M., 2008. Noise in the nervous system. Nat. Rev. Neurosci. 9, 292–303. https://doi.org/10.1038/nrn2258

Fernald, A., 1992. Human Maternal Vocalizations to Infants as Biologically Relevant SignalsL An Evolutionary Perspective, in: Barkow, J.H., Cosmides, L., Tooby, J. (Eds.), The Adapted Mind: Evolutionary Psychology and the Generation of Culture. Oxford University Press, pp. 391–428.

Fuchs, E., Ayali, A., Robinson, A., Hulata, E., Ben-Jacob, E., 2007. Coemergence of regularity and complexity during neural network development. Dev. Neurobiol. 67, 1802–1814. https://doi.org/10.1002/dneu.20557

Gamer, M., Lemon, J., Fellows Puspendra Singh, I., 2019. irr: Various Coefficients of Interrater Reliability and Agreement.

Garrett, D.D., McIntosh, A.R., Grady, C.L., 2011. Moment-to-moment signal variability in the human brain can inform models of stochastic facilitation now. Nat. Rev. Neurosci. 12, 612– 612. https://doi.org/10.1038/nrn3061-c1

Garrett, D.D., Samanez-Larkin, G.R., MacDonald, S.W.S., Lindenberger, U., McIntosh, A.R., Grady, C.L., 2013. Moment-to-moment brain signal variability: A next frontier in human brain mapping? Neurosci. Biobehav. Rev. 37, 610–624. https://doi.org/10.1016/j.neubiorev.2013.02.015

Gorgolewski, K.J., Auer, T., Calhoun, V.D., Craddock, R.C., Das, S., Duff, E.P., Flandin, G., Ghosh, S.S., Glatard, T., Halchenko, Y.O., Handwerker, D.A., Hanke, M., Keator, D., Li, X., Michael, Z., Maumet, C., Nichols, B.N., Nichols, T.E., Pellman, J., Poline, J.B., Rokem, A., Schaefer, G., Sochat, V., Triplett, W., Turner, J.A., Varoquaux, G., Poldrack, R.A., 2016. The brain imaging data structure, a format for organizing and describing outputs of neuroimaging experiments. Sci. Data 3, 1–9. https://doi.org/10.1038/sdata.2016.44

Gottlieb, G., 1971. Ontogenesis of sensory function in birds and mammals, in: Tobach, E., Aronson, L.R., Shaw, E. (Eds.), The Biopsychology of Development. Academic Press, New York, NY, pp. 67–128.

Grandy, T.H., Garrett, D.D., Schmiedek, F., Werkle-Bergner, M., 2016. On the estimation of brain signal entropy from sparse neuroimaging data. Sci. Rep. 6, 23073. https://doi.org/10.1038/srep23073

Gurau, O., Bosl, W.J., Newton, C.R., 2017. How Useful Is Electroencephalography in the Diagnosis of Autism Spectrum Disorders and the Delineation of Subtypes: A Systematic Review. Front. Psychiatry 8, 121. https://doi.org/10.3389/fpsyt.2017.00121

Hasegawa, C., Takahashi, T., Yoshimura, Y., Nobukawa, S., Ikeda, T., Saito, D.N., Kumazaki, H., Minabe, Y., Kikuchi, M., 2018. Developmental Trajectory of Infant Brain Signal Variability: A Longitudinal Pilot Study. Front. Neurosci. 12, 566. https://doi.org/10.3389/fnins.2018.00566

Hoehl, S., Wahl, S., 2012. Recording Infant ERP Data for Cognitive Research. Dev. Neuropsychol. 37, 187–209. https://doi.org/10.1080/87565641.2011.627958

Kang, J., Chen, H., Li, Xin, Li, Xiaoli, 2019. EEG entropy analysis in autistic children. J. Clin. Neurosci. 62, 199–206. https://doi.org/10.1016/J.JOCN.2018.11.027

Kang, J., Zhou, T., Han, J., Li, X., 2018. EEG-based multi-feature fusion assessment for autism. J. Clin. Neurosci. 56, 101–107.

Kaur, Y., Ouyang, G., Junge, M., Sommer, W., Liu, M., Zhou, C., Hildebrandt, A., 2019. The reliability and psychometric structure of Multi-Scale Entropy measured from EEG signals at rest and during face and object recognition tasks. J. Neurosci. Methods 326, 108343. https://doi.org/10.1016/J.JNEUMETH.2019.108343

Kosciessa, J.Q., Kloosterman, N.A., Garrett, D.D., 2020. Standard multiscale entropy reflects neural dynamics at mismatched temporal scales: What’s signal irregularity got to do with it? PLoS Comput. Biol. 16, e1007885. https://doi.org/10.1371/journal.pcbi.1007885

Kuntzelman, K., Jack Rhodes, L., Harrington, L.N., Miskovic, V., 2018. A practical comparison of algorithms for the measurement of multiscale entropy in neural time series data. Brain Cogn. 123, 126–135. https://doi.org/10.1016/J.BANDC.2018.03.010

Leach, S.C., Morales, S., Bowers, M.E., Buzzell, G.A., Debnath, R., Beall, D., Fox, N.A., 2020. Adjusting ADJUST: Optimizing the ADJUST algorithm for pediatric data using geodesic nets. Psychophysiology 57, e13566. https://doi.org/10.1111/PSYP.13566

Lippé, S., Kovacevic, N., McIntosh, A.R., 2009. Differential maturation of brain signal complexity in the human auditory and visual system. Front. Hum. Neurosci. 3, 48. https://doi.org/10.3389/neuro.09.048.2009

Liu, T., Chen, Y., Chen, D., Li, C., Qiu, Y., Wang, J., 2017. Altered electroencephalogram complexity in autistic children shown by the multiscale entropy approach. Neuroreport 28, 169–173. https://doi.org/10.1097/WNR.0000000000000724

Lopez-Calderon, J., Luck, S.J., 2014. ERPLAB: an open-source toolbox for the analysis of event-related potentials. Front. Hum. Neurosci. 8, 213. https://doi.org/10.3389/fnhum.2014.00213

Luck, S.J., 2014. An Introduction to the Event-Related Potential Technique - Steven J. Luck - Google Books, 2nd ed. MIT Press.

McIntosh, A.R., Kovacevic, N., Itier, R.J., 2008. Increased brain signal variability accompanies lower behavioral variability in development. PLoS Comput. Biol. 4. https://doi.org/10.1371/journal.pcbi.1000106

Mišić, B., Doesburg, S.M., Fatima, Z., Vidal, J., Vakorin, V.A., Taylor, M.J., McIntosh, A.R., 2015. Coordinated information generation and mental flexibility: Large-scale network disruption in children with autism. Cereb. Cortex 25, 2815–2827. https://doi.org/10.1093/cercor/bhu082

Miskovic, V., Owens, M., Kuntzelman, K., Gibb, B.E., 2016. Charting moment-to-moment brain signal variability from early to late childhood. Cortex. 83, 51–61. https://doi.org/10.1016/j.cortex.2016.07.006

Mognon, A., Jovicich, J., Bruzzone, L., Buiatti, M., 2011. ADJUST: An automatic EEG artifact detector based on the joint use of spatial and temporal features. Psychophysiology 48, 229– 240. https://doi.org/10.1111/j.1469-8986.2010.01061.x

Mumme, D.L., Fernald, A., Herrera, C., 1996. Infants’ Responses to Facial and Vocal Emotional Signals in a Social Referencing Paradigm. Child Dev. 67, 3219–3237. https://doi.org/10.1111/j.1467-8624.1996.tb01910.x

Nikulin, V. V, Brismar, T., 2004. Comment on “Multiscale Entropy Analysis of Complex Physiologic Time Series.” Phys. Rev. Lett. 92, 89803. https://doi.org/10.1103/PhysRevLett.92.089803

Nolan, H., Whelan, R., Reilly, R.B., 2010. FASTER: Fully Automated Statistical Thresholding for EEG artifact Rejection. J. Neurosci. Methods 192, 152–162. https://doi.org/10.1016/j.jneumeth.2010.07.015

Okazaki, R., Takahashi, T., Ueno, K., Takahashi, K., Ishitobi, M., Kikuchi, M., Higashima, M., Wada, Y., 2015. Changes in EEG Complexity with Electroconvulsive Therapy in a Patient with Autism Spectrum Disorders: A Multiscale Entropy Approach. Front. Hum. Neurosci. 9, 106. https://doi.org/10.3389/fnhum.2015.00106

Pernet, C.R., Appelhoff, S., Gorgolewski, K.J., Flandin, G., Phillips, C., Delorme, A., Oostenveld, R., 2019. EEG-BIDS, an extension to the brain imaging data structure for electroencephalography. Sci. Data. https://doi.org/10.1038/s41597-019-0104-8

Polizzotto, N.R., Takahashi, T., Walker, C.P., Cho, R.Y., 2016. Wide range multiscale entropy changes through development. Entropy 18, 12. https://doi.org/10.3390/e18010012

Puglia, M.H., Krol, K.M., Missana, M., Williams, C.L., Lillard, T.S., Morris, J.P., Connelly, J.J., Grossmann, T., 2020. Epigenetic tuning of brain signal entropy in emergent human social behavior. BMC Med. 18, 244. https://doi.org/10.1186/s12916-020-01683-x

R Core Team, 2020. R: A language and environment for statistical computing.

Rezaeezadeh, M., Shamekhi, S., Shamsi, M., 2020. Attention Deficit Hyperactivity Disorder Diagnosis using non-linear univariate and multivariate EEG measurements: a preliminary study. Phys. Eng. Sci. Med. 43, 577–592. https://doi.org/10.1007/s13246-020-00858-3

Richman, J.S., Moorman, J.R., 2000. Physiological time-series analysis using approximate entropy and sample entropy. Am. J. Physiol. Circ. Physiol. 278, H2039–H2049. https://doi.org/10.1152/ajpheart.2000.278.6.H2039

Sathyanarayana, A., El Atrache, R., Jackson, M., Alter, A.S., Mandl, K.D., Loddenkemper, T., Bosl, W.J., 2020. Nonlinear Analysis of Visually Normal EEGs to Differentiate Benign Childhood Epilepsy with Centrotemporal Spikes (BECTS). Sci. Rep. 10, 1–12. https://doi.org/10.1038/s41598-020-65112-y

Shafiei, G., Zeighami, Y., Clark, C.A., Coull, J.T., Nagano-Saito, A., Leyton, M., Dagher, A., Mišić, B., 2019. Dopamine Signaling Modulates the Stability and Integration of Intrinsic ć Brain Networks. Cereb. Cortex 29, 397–409. https://doi.org/10.1093/cercor/bhy264

Shew, W.L., Yang, H., Petermann, T., Roy, R., Plenz, D., 2009. Neuronal Avalanches Imply Maximum Dynamic Range in Cortical Networks at Criticality. J. Neurosci. 29, 15595–15600. https://doi.org/10.1523/JNEUROSCI.3864-09.2009

Simon, D.M., Damiano, C.R., Woynaroski, T.G., Ibañez, L. V., Murias, M., Stone, W.L., Wallace, M.T., Cascio, C.J., 2017. Neural Correlates of Sensory Hyporesponsiveness in Toddlers at High Risk for Autism Spectrum Disorder. J. Autism Dev. Disord. 47, 2710– 2722. https://doi.org/10.1007/s10803-017-3191-4

Stein, R.B., Gossen, E.R., Jones, K.E., 2005. Neuronal variability: noise or part of the signal? Nat. Rev. Neurosci. 6, 389–397. https://doi.org/10.1038/nrn1668

Szostakiwskyj, J.M.H.H., Willatt, S.E., Cortese, F., Protzner, A.B., 2017. The modulation of EEG variability between internally- and externally-driven cognitive states varies with maturation and task performance. PLoS One 12, e0181894. https://doi.org/10.1371/journal.pone.0181894

Takahashi, T., Cho, R.Y., Mizuno, T., Kikuchi, M., Murata, T., Takahashi, K., Wada, Y., 2010. Antipsychotics reverse abnormal EEG complexity in drug-naive schizophrenia: A multiscale entropy analysis. Neuroimage 51, 173–182. https://doi.org/10.1016/j.neuroimage.2010.02.009

Vakorin, V.A., McIntosh, A.R., Mišić, B., Krakovska, O., Poulsen, C., Martinu, K., Paus, T., 2013. Exploring Age-Related Changes in Dynamical Non-Stationarity in Electroencephalographic Signals during Early Adolescence. PLoS One 8, e57217. https://doi.org/10.1371/journal.pone.0057217

Wadhera, T., Kakkar, D., 2020. Conditional entropy approach to analyze cognitive dynamics in autism spectrum disorder. Neurol. Res. 869–878. https://doi.org/10.1080/01616412.2020.1788844

Walker-Andrews, A.S., 1997. Infants’ perception of expressive behaviors: Differentiation of multimodal information. Psychol. Bull. 121, 437–456. https://doi.org/10.1037/0033-2909.121.3.437

Ward, L.M., Doesburg, S.M., Kitajo, K., MacLean, S.E., Roggeveen, A.B., 2006. Neural synchrony in stochastic resonance, attention, and consciousness. Can. J. Exp. Psychol. Can. Psychol. expérimentale 60, 319.

Weng, W.C., Chang, C.F., Wong, L.C., Lin, J.H., Lee, W.T., Shieh, J.S., 2017. Altered resting-state EEG complexity in children with tourette syndrome: A preliminary study. Neuropsychology 31, 395–402. https://doi.org/10.1037/neu0000363

Williams, C.L., Puglia, M.H., 2021. APPLESEED Example Dataset. https://doi.org/10.18112/openneuro.ds003710.v1.0.0

Zhang, D., Ding, Haiyan, Liu, Y., Zhou, C., Ding, Haishu, Ye, D., 2009. Neurodevelopment in newborns: a sample entropy analysis of electroencephalogram. Physiol. Meas. 30, 491–504. https://doi.org/10.1088/0967-3334/30/5/006

